# Circular continuum of alpha motoneuron types

**DOI:** 10.1101/2025.08.12.669858

**Authors:** Teresa L. Garrett, Lee Wintermute, Maggie Armitage, Shelby Ward, Ibrahim Abdul Halim, Mohamed H. Mousa, Kalin Gerber, Sherif M. Elbasiouny

## Abstract

Alpha-motoneurons (α-MNs) are traditionally classified into slow (S), fast fatigue-resistant (FR), and fast-fatigable (FF), which exist along a continuum of properties between slow and fast, enabling the generation of graded force and seamless movement. Using combinations of markers, we developed novel immunohistochemistry protocols that enabled co-labeling of six major and transitional α-MN types throughout the mouse lumbar spinal cord with unprecedented detail. Intriguingly, our protocols labeled for the first time: α-MNs of the fast fatigue intermediate (FI) type; a previously undescribed transitional α-MN subtype (FR/FI); and a novel subtype of α-MNs exhibiting hybrid characteristics of both S and FF types – termed S/FF – which resist ALS degeneration. Electrophysiological recordings confirmed FR/FI and S/FF subtypes, both exhibiting mixed traits. The discovery of S/FF subtype reveals that α-MNs exist along a circular continuum between slow and fast types, challenging the traditional linear model and reshaping our understanding of their role in motor control.

**New & Noteworthy:** This study introduces a novel immunohistochemistry protocol to co-label six distinct types of adult spinal cord motor neurons. This technical innovation led to a conceptual paradigm shift, revealing a circular continuum of motor neuron subtypes rather than a linear one. This new framework provides unprecedented precision for studying their varying susceptibility to degeneration in diseases like ALS, with broad implications for understanding motor control in health and disease.

## INTRODUCTION

Decades of research have established that spinal alpha motor neurons (α-MNs), muscle fibers, and motor units (MUs) do not exhibit uniform functional characteristics but instead exist along a linear continuum of discrete classes ranging from slow to fast phenotypes. This gradient reflects distinct physiological and metabolic properties, including contraction speed, fatigue resistance, and force generation capacity^1,2^. Slow-twitch (Type I) muscle fibers, innervated by smaller α-MNs, demonstrate high fatigue resistance, rendering them well-suited for sustained, low-intensity motor activities^3,4^. In contrast, fast-twitch (Type II) fibers, associated with larger α-MNs, are optimized for rapid, forceful contractions but exhibit reduced fatigue resistance, making them ideal for brief, high-intensity movements. The recruitment of these MN types adheres to the size principle, wherein smaller, slower α-MNs are activated first, followed sequentially by larger, faster α-MNs as the demand for force production escalates^5^. Importantly, the physiological properties of α-MNs are closely matched to those of the muscle fibers they innervate^6^, such that slow α-MNs synapse with fatigue-resistant, slow-twitch fibers, while fast α-MNs innervate fast-twitch fibers optimized for power and speed. This hierarchical recruitment and precise matching are critical for the smooth execution of motor tasks, ensuring efficient force generation across diverse physical activities.

α-MNs can be classified based on a range of characteristics, including electrophysiological properties^7^ (such as action potential firing thresholds and afterhyperpolarization, AHP, amplitude and duration), morphological features^8,9^ (such as soma size and dendritic architecture), contractile properties (related to the speed and endurance of the muscle fibers they innervate), and immunohistochemical (IHC) markers that identify specific proteins or molecular expressions^6^. Traditionally, α-MNs have been categorized into three primary types: slow (S), fast fatigue-resistant (FR), and fast fatigable (FF), reflecting differences in their physiological roles and muscle activation profiles. Electrophysiological recordings have successfully distinguished these types, and in some cases, even identified a fourth subtype—fast intermediate (FI)^7^. However, immunofluorescence (IF) protocols have never been capable – in any species – of labeling the FI α-MN type or simultaneously co-labeling more than two α-MN types within a single sample. This significant limitation restricts comprehensive mapping of α-MN diversity and highlights the need for enhanced IF methods to fully capture the complexity of α-MN subtypes in both physiological and pathological states.

Here we developed a novel IF protocol capable of co-labeling six distinct types (4 major types and 2 transitional subtypes) of α-MNs in the lumbar spinal cord of the adult mouse, thereby enabling an unprecedented level of detail. Using this protocol, in addition to examining the distribution of different α-MN types across the lumbar cord, we identified two novel transitional subtypes of spinal α-MNs, FR/FI and S/FF, which shares molecular, electrical, and morphological features of both S and FF α-MNs. Our results show that these newly described cells are present in male and female wild-type (WT) mice and do not degenerate in ALS. Electrophysiological recordings confirmed the presence of both FR/FI and S/FF α-MNs with mixed slow and fast characteristics, a phenomenon that could only be replicated in simulations when both slow SK3 and fast SK2 currents were present. This finding indicates that α-MNs are arranged along a circular continuum, contradicting the conventional view of a linear progression from S to FF types as opposing extremes.

## RESULTS

### Typing major α-MN types, including the FI type, in the mouse spinal cord

Molecular markers, in the right combinations, can distinguish the major types of α-MNs (**Fig 1**). Our first objective was to use a combination of slow (SK3) and fast (OPN and MMP9) markers to categorize α-MNs in the adult mouse spinal cord into their respective types. In addition to these markers, we incorporated NeuN (a neuronal marker) and VaChT (a cholinergic synapse marker specific to α-MNs) to label and identify α-MN cell bodies. Therefore, we employed two distinct immunofluorescence (IF) protocols on serial sections of spinal cord tissue (protocols 1 and 2) to label α-MNs with these five markers. **Figure 1** illustrates these protocols and demonstrates how the expression patterns of these markers successfully distinguish the four major α-MN types.

**Figure 1.**
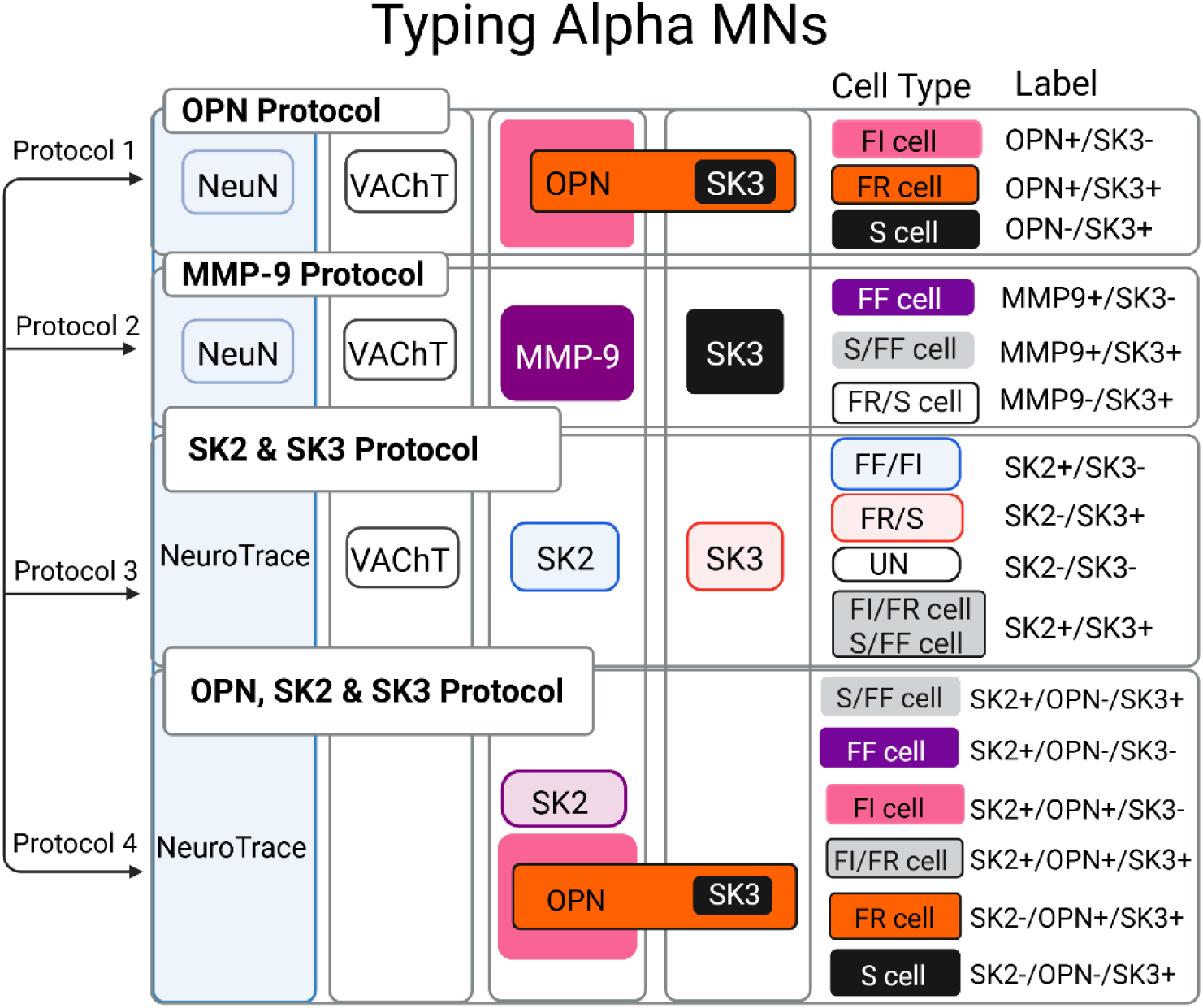
Overview of immunofluorescence protocols and antibody combinations used for α-MN typing. Each protocol is numbered and corresponds to a specific combination of antibodies, color-coded to indicate the labeled cell type. The figure shows which cell types are identified by each antibody combination. (figure created using Biorender)

To validate the IF protocols, we measured the largest cross-sectional area (LCA) of labeled α-MNs in the lumbar spinal cord of adult SOD1-G93A WT male mice (3-4 months old) to verify that our methods reproduce the known soma size differences among α-MN types. Our results show that the mean LCA of the labeled cells aligns with established size differences (**Fig. 2C**): cells expressing MMP9 but not SK3 (MMP9+/SK3–, probable FF cells) were the largest, followed by cells expressing OPN but not SK3 (OPN+/SK3–, probable FI cells), which were larger than cells co-expressing OPN and SK3 (OPN+/SK3+, probable FR cells). The smallest cells were those expressing SK3 but not OPN (OPN–/SK3+, probable S cells). These differences in mean LCA were statistically significant (p<0.001, **Fig. 2B**). Similar size differences among α-MN types were observed in females (see **Supplemental Fig. 1A**).

**Figure 2.**
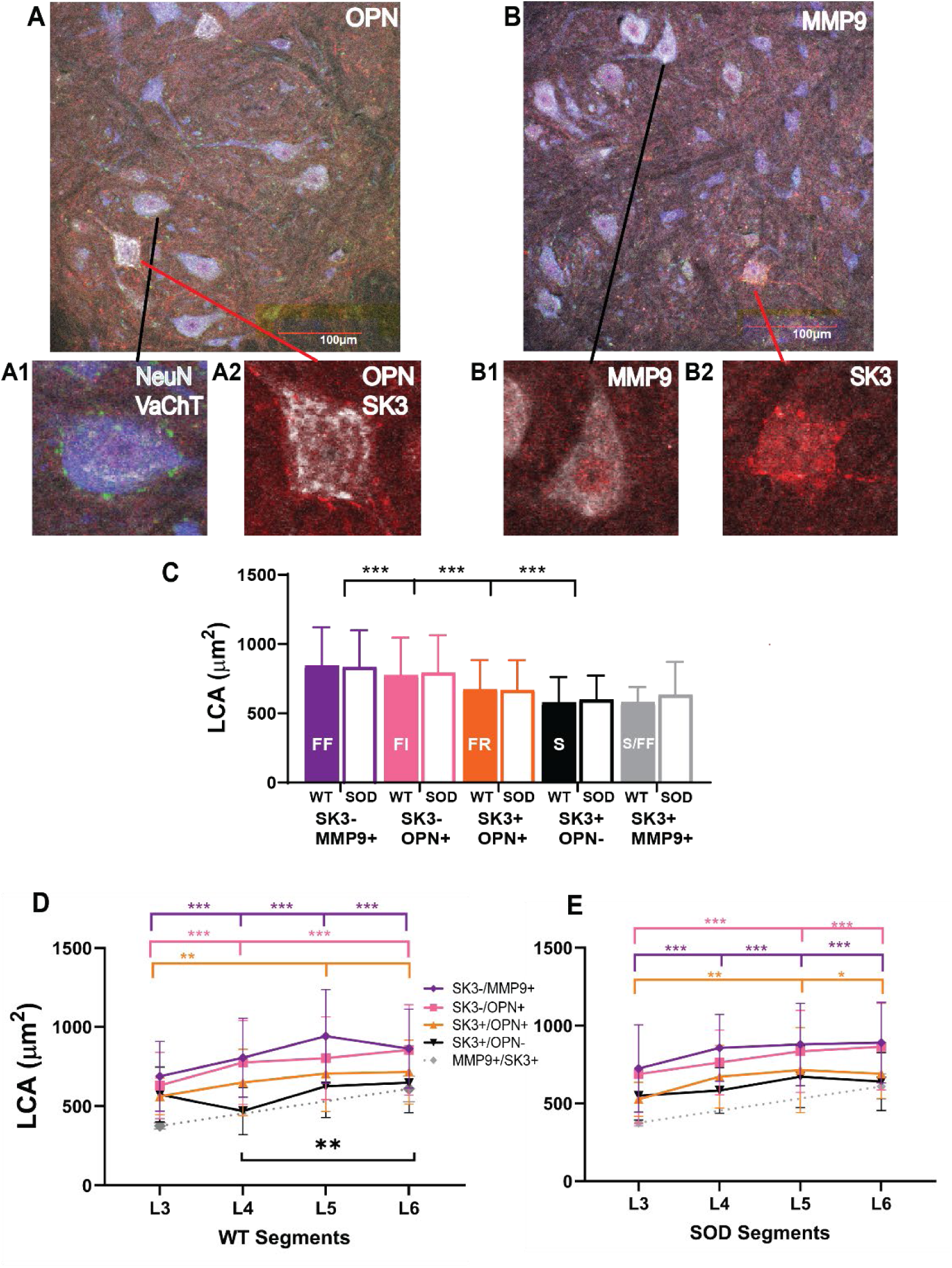
α-MN types of P90 SOD1-G93A WT and SOD (symptom onset) mice differentiated using protocols 1 and 2. (**A**) 20x image of protocol 1: OPN, SK3, NeuN and VaChT, (**A1**) Zoom of MN with NeuN (blue) and VaChT (green) and (**A2**) OPN (white) and SK3 (red) labels (**B**) 20x image of protocol 2: MMP9, SK3, NeuN and VaChT (**B1**) Zoom of cell with MMP9 (white) and (**B2**) SK3 (red) label. Black and red lines indicate where the cells are in the sections. (**C**) Mean LCA changes of α-MN types in WT (N of each type FF=1115, FI=966, FR=186=, S=83, and S/FF=16) and SOD (N of each type FF=677, FI=539, FR=133, S=117, and S/FF=19) mice. LCA changes across lumbar segments in WT (**D**) and SOD (**E**) mice for each α-MN type. Error bars represent SD. Statistical significance: *** p ≤ 0.001, ** p ≤ 0.003, * p ≤ 0.05.

To determine whether the size differences described above are consistent throughout the lumbar cord, we examined the LCA of various α-MN types across lumbar segments L3-L6. Although overall cell size generally increased from L3 to L6 (**Table S2**), the relative size differences among α-MN types remained consistent across these segments. Specifically, MMP9+/SK3– (probable FF) cells consistently exhibited the largest size, while OPN–/SK3+ (probable S) cells were the smallest. The sizes of OPN+/SK3– (probable FI) and OPN+/SK3+ (probable FR) cells fell between these extremes (**Fig. 2D**). Furthermore, these size trends were consistent in both male and female mice (see **Supplemental Fig. 1C**). Collectively, these results confirm that the IF protocols developed can reliably co-label the four major α-MN types, including the FI type, in the lumbar spinal cord of adult mice.

### MMP9+/SK3+ cells: a MN type that co-express slow and fast markers and resist ALS degeneration

Interestingly, our labeling of α-MN types revealed a small subset of lumbar MNs that co-expressed both slow and fast markers (MMP9+/SK3+). These transitional cells, which have not been previously described in the literature, are termed the “S/FF” type. Their average somatic LCA is comparable to that of S MNs (**Fig. 2C**, gray bar), suggesting that they are structurally more similar to S MNs. Notably, these cells were only observed in certain lumbar segments (specifically L3 and L6; **Fig. 2D**) and, when present, were found surrounded by other MN types. Their LCA increases from L3 to L6 (**Fig. 2D**), and among all α-MN types in these segments, S/FF cells exhibit the lowest LCA (**Fig. 2D**) and density (**Figs. 3B and 3C**). They constitute less than 0.7% of all α-MN types in the adult mouse lumbar cord (see the small gray slice in **Figs. 3D & E**). S/FF cells are also present in the lumbar cords of adult female mice (**Supplemental Fig. 1**).

**Figure 3.**
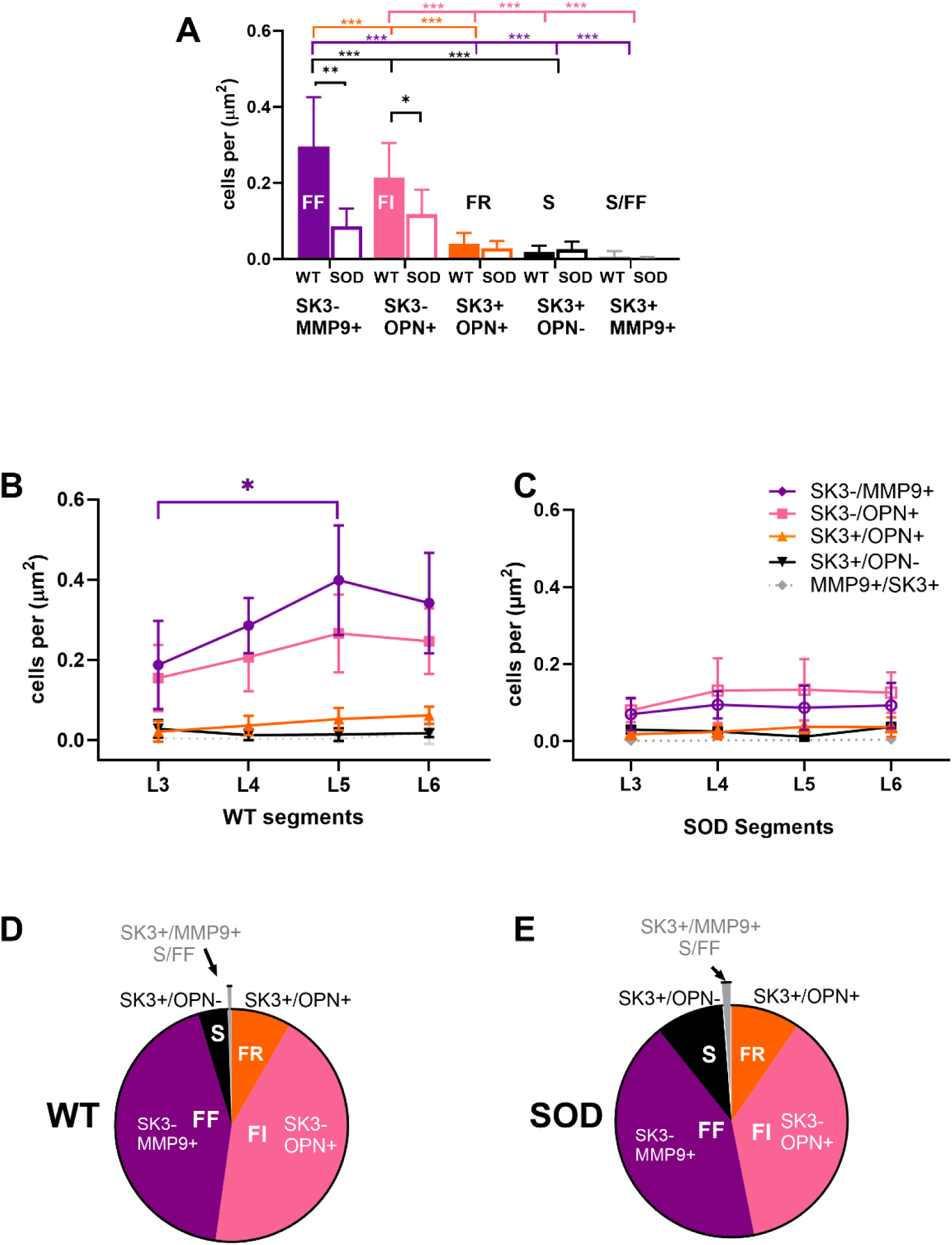
α-MN density across types using protocols 1 & 2. (**A**) Mean density changes of α-MN types in SOD1-G93A WT and SOD mice. Density changes across lumbar segments in WT (**B**) and SOD (**C**) mice for each α-MN type. Percent distribution of each α-MN type in (**D**) WT and (**E**) SOD mice. Error bars represent SD. Statistical significance: *** p ≤ 0.001, ** p ≤ 0.003, * p ≤ 0.05.

Given that transitional S/FF MNs share molecular properties with both disease-vulnerable FF MNs and disease-resistant S MNs, we examined their susceptibility to ALS degeneration. Our results show that SOD spinal cords contain comparable numbers and proportions of S/FF MNs to those in WT mice (**Figs. 3A, B, & C**), indicating that these cells are resistant to ALS degeneration. Conversely, FI MNs, as expected, had lower numbers and proportions in SOD spinal cords relative to WT, indicating their vulnerability in the disease.

### SK2 and SK3 co-labeling parse α-MNs into different, yet overlapping, types

The identification of major α-MN types in the experiments above required two separate IF protocols, resulting in distinct sets of tissue samples in which spinal motoneurons did not co-express all markers. To overcome this limitation, we developed a single IF protocol capable of labeling the four major α-MN types. Deardorff et al.^10^ demonstrated that SK2 and SK3 isoforms differentially label α-MNs—with SK3 exclusively marking slow α-MNs and SK2 labeling all α-MN types. However, their sequential IF protocol, which applied the SK3 antibody first followed by the SK2 antibody on the same tissue, may have artificially enhanced the widespread expression of SK2. To address this, we re-evaluated the expression patterns of SK2 and SK3 in α-MNs using a revised IF protocol (Protocol 3, **Fig. 1**) that enables simultaneous co-labeling by employing antibodies derived from different species for more accurate detection.

Our results show that the vast majority of α-MNs (≈92.2%) do not co-express SK2 and SK3 channels (**Fig. 4C**). As illustrated in **Fig. 4C**, approximately 70.9% of cells express SK2 only (SK3-/SK2+), ≈21.3% express SK3 only (SK3+/SK2-), ≈7.7% co-express both (SK3+/SK2+), and only ≈0.1% express neither (SK3-/SK2-). SK3-/SK2+ cells exhibited the largest somatic LCA mean (**Fig. 4B**, red bar) and distribution (**Supplemental Fig. 2C**, red trace), indicating that they comprise the fastest types (FF and FI cells). In contrast, SK3+/SK2-cells had the smallest somatic LCA mean (**Fig. 4B**, blue bar) and distribution (**Supplemental Fig. 2C**, blue trace), corresponding to the slowest types (S and FR cells). The small subset of SK3+/SK2+ cells showed an intermediate somatic LCA range (**Fig. 4B**, gray bar) and distribution (**Supplemental Fig. 2C**, gray trace), suggesting that they represent transitional groups between fast and slow cells. However, since S/FF cells have LCA similar to S cells (**Fig. 2C**) while SK3+/SK2+ cells exhibit intermediate LCAs between the fast (FF & FI cells) and slow types (FR & S cells, **Fig. 4B**), this implies the existence of an additional transitional subtype, that are larger in size bridging fast and slow cells. Additionally, the few SK3-/SK2-cells – which, despite being VaChT positive, lack SK labeling – displayed a small somatic LCA mean (similar to the FR&S group, **Figs. 4B and 4C**, white bar) and a peak distribution overlapping with intermediate types (**Supplemental Fig. 2C**, dashed trace); we have designated these as an unknown (UN) group (see Discussion). Interestingly, the four α-MN groups identified using the SK2 and SK3 co-labeling protocol, along with the UN group, were present in all lumbar segments and maintained consistent somatic LCA relationships across segments (**Fig. 4D**).

**Figure 4.**
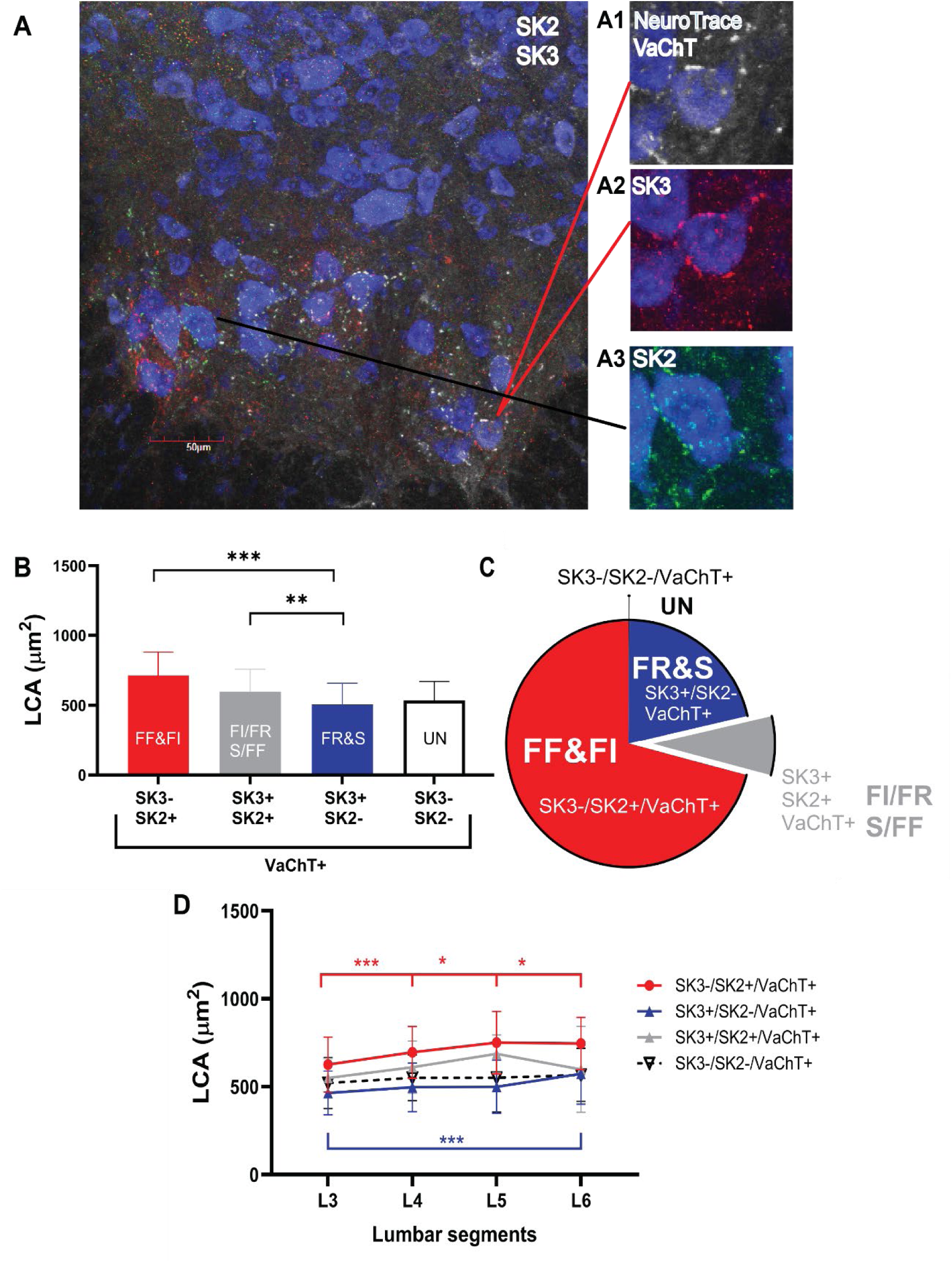
α-MN types of SOD1-G93A WT mice differentiated using protocol 3. (A) 20x image of protocol 3: OPN, SK2, SK3 and NeuroTrace (A1) Zoom cell labeled with NeuroTrace (blue), VaCht (white) and (A2) SK3 (red) (A3) SK2 (green). Black and red lines indicate where in the section the cells are located (B) LCA of α-MN types identified by SK2 and SK3 markers (protocol 3). (**C**) Percent distribution of α-MN types labeled with SK2 and SK3. (**D**) LCA variation of WT α-MN types across lumbar segments. Error bars represent SD. Statistical significance: *** p ≤ 0.001, ** p ≤ 0.003, * p ≤ 0.05.

In summary, our SK2 and SK3 co-labeling protocol demonstrates that: 1) The vast majority of α-MNs do not co-express SK2 and SK3 isoforms (with SK2 predominantly expressed by fast MNs and SK3 by slow MNs), 2) SK2 and SK3 isoforms can parse α-MNs into distinct yet overlapping groups, and 3) Although VaChT is a marker of α-MNs, SK2 and SK3 co-labeling can identify ≈99.9% of α-MNs without the need for VaChT.

### SK2, SK3, and OPN co-labeling parse α-MNs into six distinct types

Based on our finding that SK2/SK3 co-labeling without VaChT can identify ≈99.9% of α-MNs, we developed an additional four-channel IHC protocol that combines SK2/SK3 co-labeling with OPN in place of VaChT (Protocol 4 in **Fig. 1**). With cells co-expressing all markers, we were able to distinguish more α-MN types than with the single protocols used previously. This protocol was tested in a different set of animals than those used in **Fig. 2**.

Using this protocol, we distinctly labeled the four major α-MN types (FF, FI, FR, and S) as well as two transitional subtypes (FI/FR and S/FF) that represent a small percentage of the population. Specifically, OPN separated SK3+/SK2-cells into the S and FR types, SK3-/SK2+ cells into the FF and FI types, and SK3+/SK2+ cells into transitional FI/FR and S/FF subtypes (**Fig. 5B**). Notably, this provides a second independent line of evidence for a transitional population of SK3+/SK2+/OPN-cells existing between S and FF α-MNs (i.e., the S/FF type), with the first evidence being the SK3+/MMP9+ cells in **Fig. 2**. The SK3+/SK2+/OPN-cells have somatic LCA means (**Fig. 5B**) between FF and S cells, consistent with a transitional S/FF subtype. Likewise, SK3+/SK2+/OPN+ cells exhibited somatic LCA values (**Fig. 5B**) intermediate between FR and FI cells, supporting their classification as transitional FR/FI subtype. Importantly, the average somatic LCA values and distributions of the S, FR, FI, and FF types identified via the SK3/SK2/OPN protocol (Protocol 4, **Fig. 5**) closely resembled those obtained with the OPN and MMP9 protocols (Protocols 1 and 2, **Fig. 2**). **Supplemental Fig. 2** provides a cross-comparison of the somatic LCA distributions from all protocols. Because the SK3/SK2/OPN protocol parsed α-MNs into more distinct groups than the OPN and MMP9 protocols, the somatic LCA distributions for some types (e.g., S and FI in **Supplemental Fig. 2D**) displayed narrower peaks confined to more specific ranges. This improved separation among α-MNs: somatic LCA values below 500µm² were predominantly S MNs, values between 500–700µm² were predominantly FI MNs, and values of 1,000µm² or greater were predominantly FF MNs.

**Figure 5.**
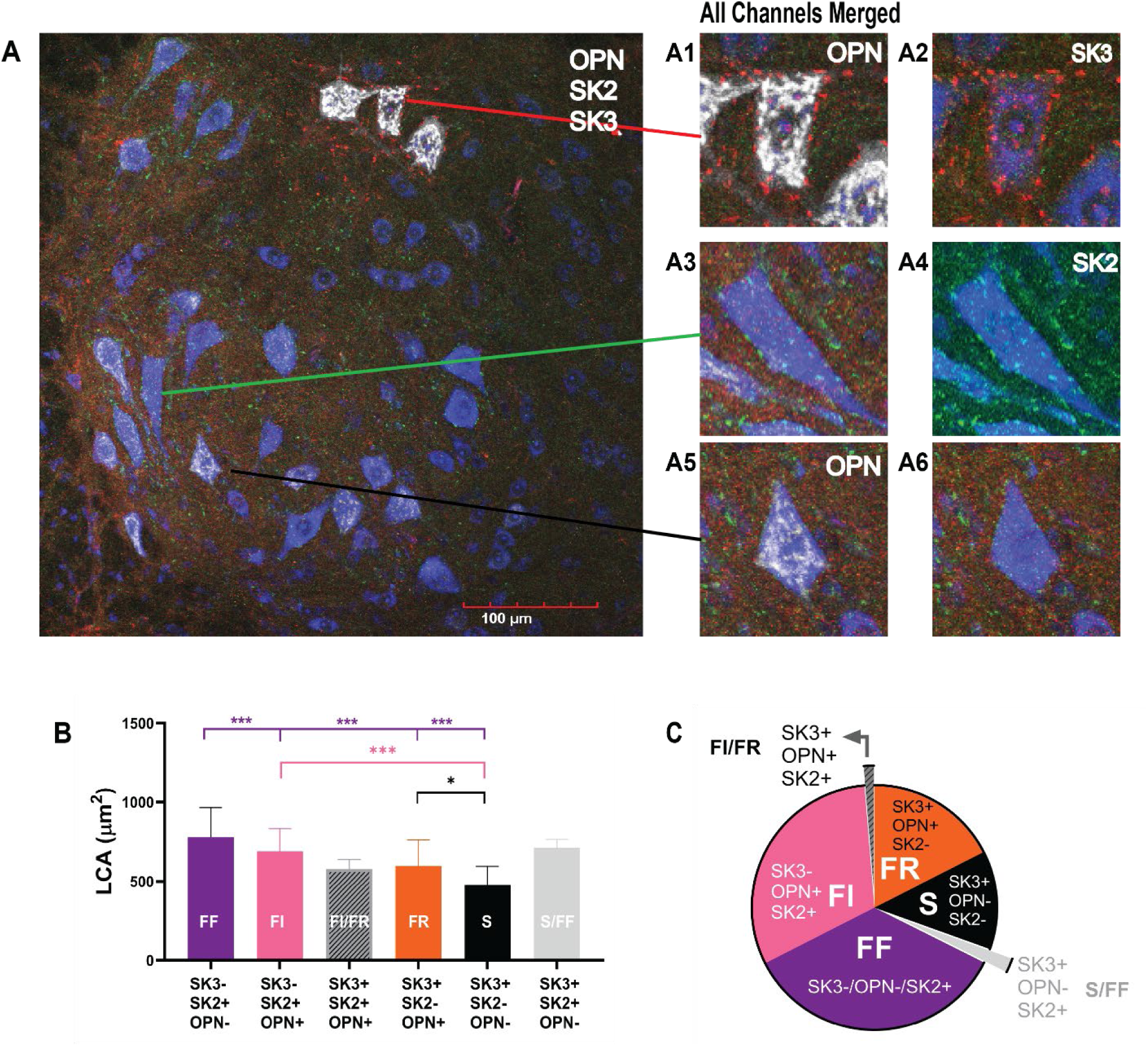
α-MN types of SOD1-G93A WT mice differentiated using protocol 4. (A) 20x image of protocol 4: OPN, SK2, SK3 and NeuroTrace (A1) Zoom of cell labeled with OPN and (A2) SK3, a cell positive SK2 (A4) and negative for A (B) LCA of α-MN types identified by SK2 and SK3 markers (protocol 3). (**B**) Percent distribution of α-MN types labeled with SK2 and SK3. (**C**) LCA variation of WT α-MN types across lumbar segments. (**D**) LCA of α-MN types identified by SK2, SK3, and OPN markers (protocol 4). (**E**) Percent distribution of α-MN types based on SK2/SK3/OPN combinations (protocol 4). Error bars represent SD. Statistical significance: *** p ≤ 0.001, * p ≤ 0.05.

In summary, the SK3/SK2/OPN protocol is a single IHC method that, through co-expression of all markers, successfully distinguishes both major and transitional α-MN types (**Fig. 5B**), further confirming the labeling observed in **Figs. 2 and 4B**. Additionally, this protocol independently confirms the existence of the transitional S/FF and FR/FI subtypes.

### Circular continuum of α-MN types

Traditionally, α-MNs have been described as existing along a linear continuum, with S-type at one extreme and FF-type at the other (**Fig. 6A**). However, the discovery of cells that co-express both slow and fast markers (S/FF and FI/FR subtypes) suggest a more complex, circular continuum of α-MN types (**Fig. 6B**). This section examines whether empirical evidence supports the existence of S/FF and FI/FR cells and a circular continuum of α-MN types.

**Figure 6.**
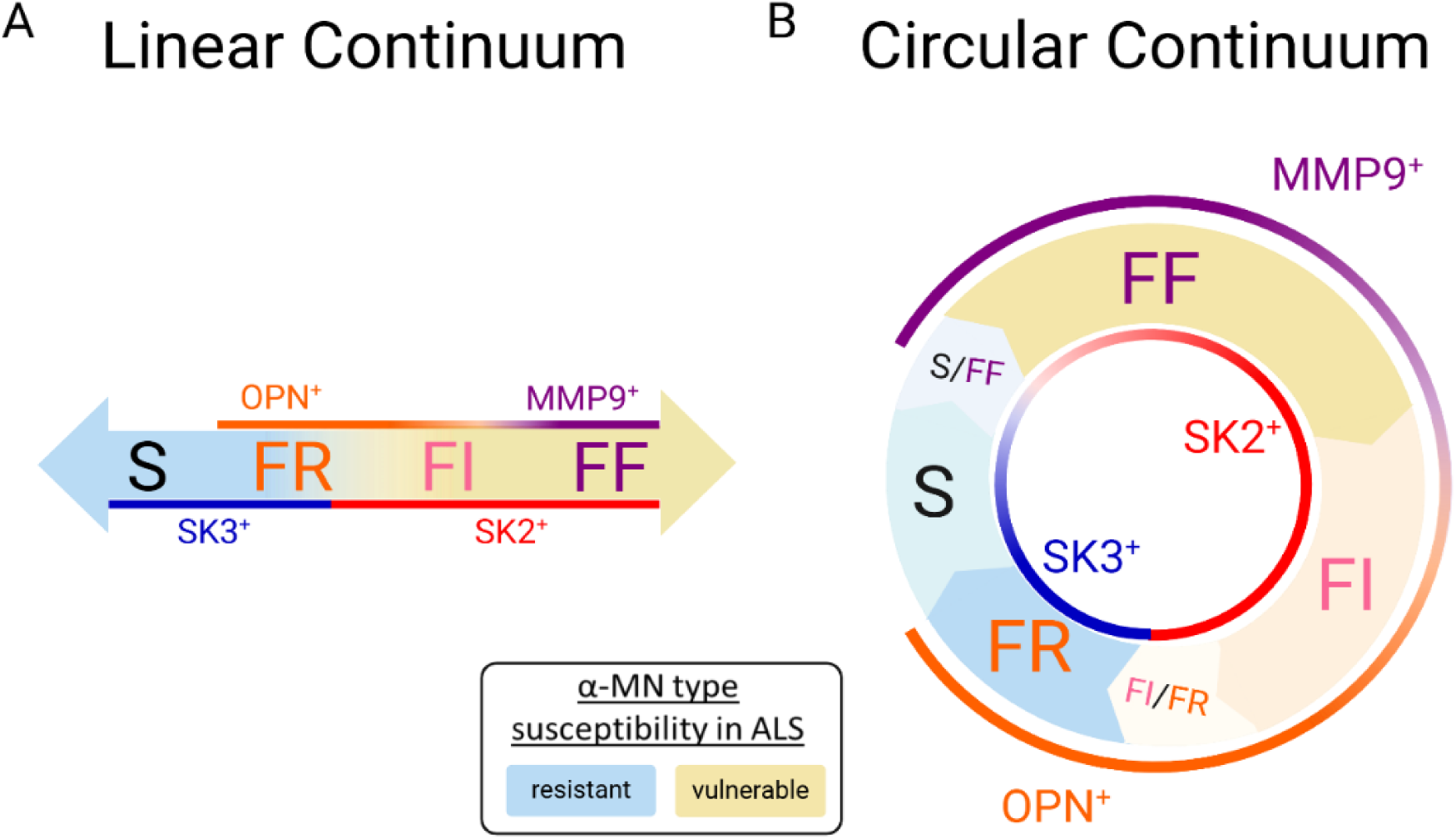
Comparison of the currently accepted linear α-MN continuum (A) with the proposed circular continuum of (B) α-MN types (figure created using Biorender).

Because α-MN types differ in their firing capacities—FF cells fire the fastest and S cells the slowest, with other types in between—and because SK channels mediate the AHP depth and duration in spinal α-MNs (with S cells exhibiting the slowest/deepest AHP and FF cells the fastest/shallowest), AHP duration serves as the primary electrophysiological property that varies with α-MN type. It can accurately classify α-MNs into their respective types^11–14^. Specifically, AHP ½ decay alone has been shown to classify α-MNs into slow and fast types with 100% accuracy— fast cells have an AHP ½ decay of less than 20ms, while slow cells have an AHP ½ decay greater than 20ms^12^. Therefore, if S/FF and FI/FR cells exist, they would be expected to exhibit hybrid slow and fast AHP dynamics, as both subtypes have been shown to co-express SK2 and SK3.

To investigate this, we obtained intracellular electrophysiological recordings from α-MNs of adult non-transgenic C57 mice and plotted the AHP amplitude against AHP ½ decay (**Fig. 7A**), with fast cells (AHP ½ decay <20ms) shown in blue and slow cells (AHP ½ decay >20ms) in red. Our IF labeling protocols indicated that SK2 channels are predominantly present in fast cells, whereas SK3 channels are predominantly found in slow cells. Therefore, we used empirical AHP recordings from a very slow cell (AHP ½ decay ≈35ms, **Figs. 7A and 7B**, outlined in green) likely to have SK3 channels only to develop a computer model of the SK3 current (red trace in **Fig. 7B** and **Table S1**), and recordings from a very fast cell (AHP ½ decay ≈10ms, **Figs. 7A and 7D**, outlined in green) likely to have to have SK2 channels only to develop a computer model of the SK2 current (blue trace in **Figs. 7B** and **Table S1**. In simulations, we optimized the SK2 and SK3 channel parameters until the AHP amplitude and duration matched the empirical data (see raw vs. simulated traces in **Figs. 7B and 7D**).

**Figure 7.**
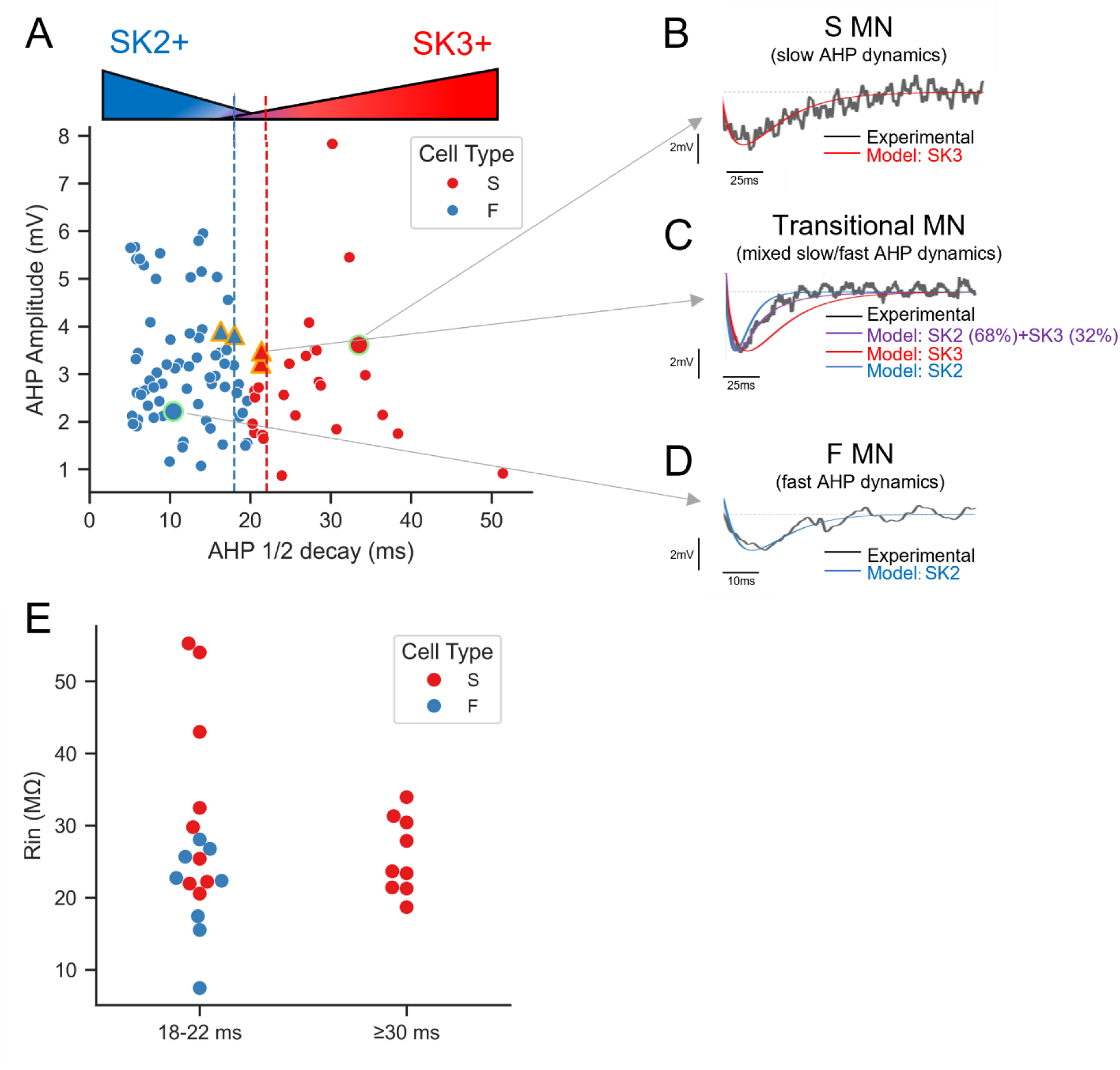
Electrophysiological recordings of MNs support a circular continuum of different α-MN types. (**A**) Empirical data of AHP amplitude vs. AHP ½ decay in non-transgenic C57 MNs (N = 99) reveal overlapping cell properties among α-MN types. The vertical blue and red dashed lines mark the range where transitional subtypes are found. Cells highlighted in yellow and green are those modeled in panels B, C, and D (indicated by gray arrows). The cells that were modeled for C (S/FF and FI/FR-type) are indicated by triangles in panel A. (**B–D**) The raw and simulated AHP recordings are shown for a S MN (**B**), a S/FF MN (**C**), and a F MN (**D**). Only SK3 (in B) and SK2 (in D) conductances could reproduce the AHP recordings of slow and fast cells, respectively. In C, both SK2 and SK3 conductances were needed to match the AHP recordings (purple trace), as SK2 alone (blue trace) or SK3 alone (red trace) did not suffice. (**E**) Comparison of Rin between transitional MNs (AHP ½ decay: 18–22ms) and the slowest slow MNs (AHP ½ decay ≥30ms) in non-transgenic C57 MNs.

Notably, the SK2 and SK3 channel models exhibited drastically different current dynamics, indicating that currents mediated by these isoforms are not merely scaled versions of each other. The SK2 channel model had a Hill coefficient (*n*_*S*2_) of 4, a half-maximal effective concentration (*K*_0.5_) of 0.5µM, and a time constant (*τ*_*S*2_) of 7ms, while the SK3 channel model had an *n*_*S*3_ of 1.3, *K*_0.5_ of 0.5µM, and *τ*_*S*3_ of 25ms. This suggests that SK3 currents activate more slowly, have a lower activation slope, and are more sensitive to Ca²⁺ compared to SK2 channels. Using these channel models, we were able to simulate the AHP recordings of most fast (blue) cells with SK2 channels alone, adjusting the channel conductance to match AHP amplitude. Similarly, most slow (red) cell recordings were accurately simulated using SK3 channels alone. However, most cells with AHP ½ decay values between 18ms and 22ms (± 2ms around the 20ms cutoff, **Fig. 7A**) could not be simulated with a single channel isoform. Instead, a combination of SK2 and SK3 currents was necessary. For example, one cell’s AHP properties were best replicated by a mix of 68% SK2 and 32% SK3 (purple trace in **Fig. 7C**). Four additional examples of cells requiring a combination of both currents are shown by the triangles in **Fig. 7A**, outlined in orange.

Because transitional S/FF and FR/FI subtypes co-express SK2 and SK3 (**Figs. 2, 3, 4, & 5**), and electrophysiological recordings suggest the presence of cells with hybrid slow and fast dynamics within the 18–22 ms AHP ½-decay range, these transitional subtypes are expected to fall predominantly within that window. As S/FF and FR/FI cells differ in size, we used input resistance (Rin) as a proxy to determine whether α-MNs within the 18–22 ms AHP ½-decay range represent size-diverse populations. For comparison, Rin values from the slowest slow-type cells (AHP ½-decay ≥30 ms, presumed S cells) were also included to indicate typical Rin values for S cells (**Fig. 7E**). Our results show that cells in the 18–22 ms range span a broad spectrum of Rin values, indicating two distinct, heterogenous populations: 1) small cell with large Rin and long AHP ½ decay>20ms), which are even smaller than the smallest S cells in the ≥30 ms group (red cells in **Fig. 7E**); and 2) large cells with low Rin and fast (of AHP ½ decay<20ms) cells (blue cells in **Fig. 7E**). This confirms our IF data that two cell populations (transitional S/FF and FR/FI subtypes) of different sizes exist in the 18–22 ms AHP ½-decay range were combined SK2 and SK3 currents mediate the AHP.

Together, these findings support our IF labeling data, confirming the existence of two transitional α-MN subtypes. These subtypes exhibit mixed slow and fast AHP dynamics – due to combined SK2 and SK3 currents – but differ markedly in cell size. These finding supports the concept of a circular continuum in α-MN types through the presence of S/FF and FI/FR transitional cells co-expressing both SK2 and SK3 channel isoforms.

## DISCUSSION

The present study identifies two novel transitional subtypes of spinal α-MNs, FR/FI and S/FF, which share molecular, electrical, and morphological features of slow and fast cells. Labeling of these cells was observed in the lumbar spinal cord of male, female, transgenic, and non-transgenic adult mice. Both electrophysiological recordings and IF staining confirmed the presence of these transitional subtypes with mixed slow and fast characteristics, a phenomenon that could only be replicated in simulations when both slow SK3 and fast SK2 currents were present. This finding demonstrates that α-MNs lie along a circular continuum, challenging the traditional view of a strictly linear progression from S to FF types as polar opposites. In addition to identifying these new subtypes, this study is the first to label the FI type. The discovery was made possible by a novel IF protocol capable of co-labeling six distinct types (4 major types and 2 transitional subtypes) of α-MNs, thereby enabling an unprecedented level of detail in studying α-MNs. This advancement opens the door to enhanced aging and neurodegenerative research where different α-MN types exhibit varying degrees of susceptibility.

For over 60 years, spinal α-MN, muscle fiber, and MU types have been extensively studied using various techniques, unveiling a continuum of morphological, biophysical, electrical, molecular, and contractile properties. Traditionally, α-MNs have been categorized along a linear spectrum between S and FF phenotypes. Electrophysiological intracellular recordings have identified up to four distinct α-MN types^7^, while immunohistochemistry (IHC) staining has differentiated up to four MU types^15^ and as many as eight muscle fiber types^16^ types. Until now, however, distinguishing α-MN subtypes using IHC has been limited. Recent RNA sequencing studies support our findings, revealing that α-MNs are far more diverse than previously recognized, with over eight distinct types identified^17,18^. Interestingly, despite these advancements, IHC staining of α-MNs have traditionally been restricted to co-labeling no more than two types simultaneously in any species^19–21^. Furthermore, no IHC protocol to date has successfully identified the FI α-MN subtype in any species.

The present study introduces a groundbreaking IF protocol that enables the co-labeling of six distinct α-MN types (FF, FI, FI/FR, FR, S, and S/FF) within the mouse lumbar spinal cord. This advancement is based on two key findings. First, we refined the understanding of SK2 and SK3 channel distribution in α-MNs, leveraging the finding that α-MNs exhibit a distinct and divergent expression pattern of SK channels. Consistent with the findings of Deardorff, et al. ^10^, our results confirm and extend that SK3 channels are exclusively expressed in slow MNs (S and FR types). However, we further demonstrate that SK2 channels are specifically expressed in fast MNs (FI and FF types), a distinction made possible by employing newer SK2 and SK3 antibodies from different species, allowing for simultaneous co-labeling of both channel types. This improvement over the sequential IF protocols used by Deardorff, et al. ^10^, reduces the risk of overestimating SK2 distribution. Second, we found that SK2 and SK3 co-labeling identifies ≈99.9% of α-MNs, effectively replacing the need for cholinergic labeling to accurately label α-MNs. Interestingly, a very small subset of α-MNs (∼0.1%) did not express either SK2 or SK3 channels. Furthermore, across all tested animals (male, female, transgenic, and non-transgenic), no cells were found that were negative for VaChT but positive for SK2 or SK3 (VaChT–, SK2+ or SK3+). This breakthrough significantly enhances the accuracy of α-MN classification and opens new avenues for detailed analysis of spinal motor circuitry.

Among all ion channels in α-MNs, SK channels are the only ones that correlate with α-MN types through their AHP properties, and AHP is the only electrical property that differ among α-MN types. Consequently, it is not surprising that the expression levels of these channel isoforms can distinguish between different α-MN types. Our findings demonstrate that SK2 and SK3 currents are not merely scaled versions of one another but exhibit markedly different current dynamics. Specifically, SK2 currents activate significantly faster than SK3 currents, with a time constant approximately 3.5 times shorter in our simulations (*τ*_*SK*2_ ≈7ms vs. *τ*_*SK*3_ ≈25ms). This difference indicates that SK2 channels not only activate more rapidly but also have a steeper activation slope and reduced sensitivity to Ca^2+^ compared to SK3 channels. These modeling data support the differential expression of these isoforms among α-MN types, highlighting their distinct functional roles. Although various markers have been used to label different α-MN types, such as SV2A^22^, Esrrb^23^, and UCHL1^24^ for slow cells; Chondrolectin^25^, CGRP ^26^, and Dlk1^27^ for fast cells; MMP9^19^ for FF cells; and Osteopontin^20^ for S and FR cells, SK2 and SK3^10^ – as ion channels that mediate the AHP they stand out as the only markers with both molecular and functional relevance, thereby providing a more accurate means of distinguishing between α-MN subtypes.

Of special interest is the finding of a transitional cell subtype, the S/FF subtype, that closes the loop between S and FF α-MNs and form a circular continuum. Our results indicate that S/FF cells are: 1) Present in both male and female mice, as well as in multiple mouse strains, 2) Small in size, with LCA and Rin values comparable to those of S cells, 3) Rare in the Lumbar cord, constituting approximately 0.7% of all MN types, and localized specifically to L3 and L6 segments, 4) Characterized by co-expression of both slow and fast markers (MMP9+/SK2+/SK3+/OPN-), 5) Exhibiting an AHP ½ decay time between 18ms and 22ms, supporting a firing rate much faster than typical S cells, and 6) Resistant to degeneration in ALS. Although these properties suggest they may represent a “super type,” it remains to be determined whether S/FF cells are fatigable or fatigue-resistant. Another important finding of this study is the successful labeling of FI α-MNs using IHC. We showed that FI cells: (1) are the second most prevalent α-MN type (after FF cells) in the adult mouse lumbar spinal cord; (2) are susceptible to degeneration in ALS; and (3) display similar size and frequency in both male and female mice. The detailed characterization of these cell types will advance research on aging and neurodegeneration, where α-MN types differ in their susceptibility.

## Material and Methods

### Immunohistochemistry

#### Ethical approval

All procedures were approved through the Wright State University animal care and use committee.

#### Animals

##### ALS mice

Mice for the immunohistochemistry studies were (∼3 months, G93A-SOD1: N = 19, WT: N = 13) bred from breeders purchased from Jackson Laboratory (B6SJL-Tg, Stock# 002726). Hybrid females were bred with hemizygote males, allowing for the expression of the SOD-1 human gene in some their offspring (SOD-1 mice) while also providing offspring that did not show this expression to be used as wildtype (WT) littermate controls. We used only male mice, as the disease is more aggressive in males^28^, to provide more consistency of disease changes and, thereby, robustness of results. Genotyping was performed by Transnetyx, Inc. Mice were randomly assigned from the colony to the study.

##### C57BL/6 mice

Mice in the supplementary figure 2 are C57BL/6, obtained from National Institute on Aging (NIA) colony (N = 9, male = 5, female = 4). The mice were randomly selected from NIA’s colony.

#### Spinal Cord Preparation

Mice were given a lethal dose of Euthasol followed by transcardial perfusion with a phosphate buffer rinse, then fixed with 4% paraformaldehyde. The spinal cords were dissected and post-fixed for 1-2hrs, then refrigerated in 15% sucrose for up to 2 weeks. Next, each segment was identified and marked with a tissue dye, embedded in tissue freezing medium, then flash-frozen and stored at −80°C. The cords were then sectioned at 45µm on a Leica cryostat (Model #: HM 550 ThermoFisher® Cryostat), serially collected, and kept in cryoprotectant at −20°C until labeling.

#### Immuno Labeling Protocols 1, 2 and 3

For the first three protocols this is the general protocol, serial slices (4-5) of each spinal cord segment were extracted from the cryoprotectant and labeled for each animal. Sections were washed three times (3x) in phosphate buffer saline with 0.1% Triton-X (PBS-T); then incubated one hour in 10% normal horse serum (NHS). The primary antibodies (see Table 1 for antibody list) were incubated overnight at 4°C; then washed 3x in PBS-T. The secondary antibodies were incubated at 25°C for 2-3 hours. After washing with phosphate buffer saline (PBS), the sections were mounted and cover-slipped with Vectashield (Vector labs) for imaging (see methods in Imaging section, below). Figures 2A, 2B and 4A, A1-A3 show examples of labeling of all protocols.

**Table 1.** List of all antibodies used in each labeling protocol with associated protocol and dilution.

| Protocol # | Antibody | Vendor | Catalog # | Dilution | RRID |
| --- | --- | --- | --- | --- | --- |
| **Primary Antibodies** |  |  |  |  |  |
| 1 & 2 | NeuN $\alpha$ guinea pig | Sigma | ABN90 | 1:300 | RRID:AB_11205592 |
| 1 & 2 | VACHT $\alpha$ mouse | Abcam | AB134298 | 1:1000 | discontinued |
| 3 | VACHT $\alpha$ mouse | Novus | NBPS-59378 | 1:400 | RRID:AB_2922997 |
| 1, 2, 3 & 4 | SK3 $\alpha$ rabbit | Sigma | AB5350 | 1:1000 | RRID:AB_3713098 |
| 1 & 4 | OPN $\alpha$ goat | R&D systems | AF808 | 1:750 | RRID:AB_2194992 |
| 2 | MMP-9 $\alpha$ goat | Sigma | M9570 | 1:500 | RRID:AB_1079397 |
| 3 & 4 | SK2 $\alpha$ goat | Abcam | AB99457 | 1:1000 | RRID:AB_10675678 |
| **Secondary Antibodies** |  |  |  |  |  |
| 1 & 2 | Alexa Fluor 405 | Jackson Immuno | 706-475-148 | 1:200 | RRID:AB_2340470 |
| 1 & 2 | Alexa Fluor 488 | Jackson Immuno | 715-545-150 | 1:100 | RRID:AB_2340846 |
| 3 | Alexa Fluor 488 | Jackson Immuno | 705-545-147 | 1:100 | RRID:AB_2336933 |
| 1, 2, 3 & 4 | Cy-3 | Jackson Immuno | 711-165-152 | 1:100 | RRID:AB_2307443 |
| 1 & 2 | Alexa Fluor 647 | Jackson Immuno | 705-605-147 | 1:100 | RRID:AB_2340437 |
| 3 & 4 | Alexa Fluor 647 | Jackson Immuno | 715-605-150 | 1:100 | RRID:AB_2340865 |
| **Other Agents** |  |  |  |  |  |
| 4 | FAB fragment | Abcam | ab98826 | 1:200 | RRID:AB_10674476 |
| 3 & 4 | Neurotrace | Life Sciences Tech | N21479 | 1:250 |  |

#### SK2, SK3 and OPN labeling (Protocol 4)

This protocol was completed in 3 SOD1-G93A WT male mice and 9 C57BL/6 male and female mice. In the SOD1-G93A WT sections were labeled with SK2, SK3, OPN, and NeuroTrace. Spinal cord segments from L5 and L6 were extracted from cryoprotectant and labeled for each animal. For the C57BL/6 mice, sections were labeled with SK2, SK3, and OPN. Spinal cord segments L3-L6 were used to complete analysis.

The SK2 was incubated overnight at 4°C; then was washed 3x with PBS-T. Then, the secondary antibody incubated for 2 hours; then FAB fragment incubated for 30min. SK3 and OPN antibodies (NeuN, same as protocol 1, was added for C57) were then incubated overnight at 4°C and, secondary antibodies incubated at 25°C for 2-3 hours; Neurotrace, (omitted for C57) was incubated for 30 min. After washing with PBS, sections were mounted and cover-slipped with Vectashield for imaging. See figure 5A & A1-A6 for example of labeling.

#### Imaging and Analysis

##### Imaging

An Olympus FV1000 confocal microscope was used to acquire 20X images at 1µm steps and 1.2x zoom of the ventral side of the spinal cord, specifically the areas containing motor neuron pools was imaged (Rexed lamina IX). Lumbar regions L3-L6 were imaged with 3-4 complete slices imaged for each lumbar segment for each animal.

##### Image Analysis

The images collected were analyzed by a blinded experimenter using Fluoview software (Olympus). The largest cross-sectional area (LCA) was measured, as well as the average immunofluorescent (IF) intensities for each cell. In protocols 1-3, cells were labeled as alpha motor neurons if they were positive for VAChT and specifically displaying c-boutons and NeuN (see **Fig 1**). Additionally, they had to have a cross-sectional area larger than 300µm (29). To determine whether cells were positive for OPN or MMP-9, background fluorescence levels were taken in an area that was negative for all markers, and the background fluorescence was averaged. Next, (i.e., OPN or MMP9), the average IF intensity of the cell at the LCA was divided against the background. If the intensity ratio was twice as high as the background, the motor neurons were categorized as positive for either MMP9 or OPN. They were then categorized into FF-, FI-, FR-, or S-motor neurons (see Fig 1). The SK2/SK3-labeled cells were determined to be positive if clusters of channels were found on the cell membrane of the soma; and were considered an α-MN if VaChT was also present. If the cell was positive for SK2 then it was considered an F (FF or FI) cell. If it was positive for SK3 was categorized as an S cell (FR or S). Labeling with OPN, SK2, and SK3 allowed us to differentiate FR (OPN+ and SK3+) and S (OPN- and SK3+) cells together with labeling of FF (OPN- and SK2+) and FI (OPN+ and SK2+) cells.

Exclusion criteria: Mice and/or images were excluded due to poor signal to noise ratios.

### Electrophysiology

#### Animals

##### C57BL/6 mice

Mice for the electrophysiology studies were both male and female C57BL/6 mice from 3 age groups were received from National Institute on Aging. The 3 age groups consisted of: 3-4 months (female N = 12 and male N = 10), 12-14 months (female N = 12 and male N = 5) and 24+ months (female N = 21 and male N = 13). The mice with the same age and sex were housed with their similar littermates, with food and water ad libitum.

#### Tissue Preparation

Mouse laminectomy of the sacrocadual region of the spinal cord was performed as previously described in Mahrous and Elbasiouny ^30,31^. Mice underwent deep anesthesia via an intraperitoneal injection of urethane (0.27g/100g weight). Animal response was checked by tail/toe pinch to ensure mice were fully anesthetized. If necessary, supplemental injections were given until the animal was unresponsive to the tail/toe pinch and respiration was observably slow and steady.

Next, the animal was securely pinned dorsal side up to a dissecting dish and supplied with a steady flow of carbogen (mixture of 95% O_2_ and 5% CO_2_) through a face mask. Afterwards, the dorsal skin was cut and removed exposing the back muscles. Longitudinal incisions were made through the back muscle down both sides of the vertebral column. Starting around the upper lumbar region (L1), a single cut to the vertebrae was made to expose the spinal cord. Once exposed, modified Artificial Cerebral Spinal Fluid (mACSF) aerated with carbogen was steadily dripped on the surgical site at a rate of 4-5 mL/min. The vertebral column was then carefully cut along the sides of the spinal cord towards the tail of the animal as far as possible to expose the sacrocaudal roots. Immediately, the spinal cord was transected, and the dura was longitudinally cut. The cord was then gently lifted to cut the ends of the spinal cord roots. Afterwards, the spinal cord was carefully transported to a mACSF filled petri dish with carbogen aeration and the animal was decapitated (an AMVA-approved method of euthanasia). At the petri dish the spinal roots were separated and freed from the spinal cord and neighboring roots. The freed cord was transected around the L6-S1 region and the S1-C02 segments were transferred to an intracellular recording chamber. Carbogen aerated normal Artificial Cerebral Spinal Fluid (nACSF) was supplied at a rate of 2.5-3 mL/min to the recording chamber. At the recording chamber the spinal cord was pinned ventral side up and the freed spinal roots were placed on bipolar electrodes. The roots were covered with petroleum jelly for cord stability and root moisture. After completing this process, the spinal cord was incubated for at least 1 hour in the recording chamber before starting intracellular recordings.

#### Solutions

The modified Artificial Cerebral Spinal Fluid (mACSF) contained the following (in millimolar): 118 NaCl, 3 KCl, 1.3 MgSO_4_, 5 MgCl_2_, 1.4 NaH_2_PO_4_, 1.5 CaCl_2_, 24 NaHCO_3_, and 25 glucose. The normal Artificial Cerebral Spinal Fluid (nACSF) solution consisted of (in millimolar): 128 NaCl, 3 KCl, 1.5 MgSO_4_, 1 NaH_2_PO_4_, 2.5 CaCl_2_, 22 NaHCO_3_, and 12 glucose. Glucose and NaHCO_3_ were added on the day of the experiment to preserve the stocks for a long period. The composition of both ACSF solutions was adopted from previous methodology (Deutsch and Elbasiouny 2024).

#### Motoneuron Identification

Sharp glass micropipettes were pulled from 2mm outer-diameter thick-walled borosilicate capillary glass (A-M Systems, Sequim, WA, USA, 604000) using a micropipette puller (P-97; Sutter Instrument, Novato, CA). Micropipettes were then filled with 3 M KC_2_H_3_O_2_ and 100 mM KCl. A silver chloride (Ag-AgCl) electrode was placed inside the micropipette for αMN recording. Micropipette resistance typically ranged from 32-43 MΩ. Micropipettes were advanced vertically into the ventral horn of the spinal cord using a micro positioner (2660; David Kopf Instruments, Tujunga, CA) until an αMN was impaled. αMNs were identified using an antidromic action potential through the stimulation of the mounted ventral roots. Stable αMNs were verified by a resting membrane potential ≤-55 mV and ≥50 mV antidromic action potential spike height. All real-time αMN recordings were interfaced using a Power 1401−3A board (Cambridge Electronic Design, Cambridge, UK) after signal amplification using an Axoclamp-2A/B amplifier (Molecular Devices, San Jose, CA, USA). The Axoclamp-2A/B was running in discontinuous current clamp (DCC) mode with 6-8 kHz switching rate. The Spike2 software (version 8.23, Cambridge Electronic Design) was utilized for data recording and storage for offline data analysis.

#### Measurement of Afterhyperpolarization Properties

The MN After-Hyperpolarization Phase (AHP) was measured by stimulating the impaled MN with 5 consecutive pules of current with 1 second intervals. The stimulated action potentials were averaged and analyzed to measure AHP amplitude, AHP 1/2 decay and AHP 2/3 decay. AHP amplitude was measured by taking the voltage difference between the resting membrane potential and the deepest voltage value of the AHP. AHP 1/2 decay was recorded as the time from the start of the AHP to the point at which the AHP amplitude had decayed to half of its value. The time at which the AHP was two thirds of its maximum value in reference to the start of the action potential was recorded as AHP 2/3 decay.

Exclusion criteria: Cells were excluded from analysis if all parameters such as AHP and resistance.

### Computational Modeling

To reverse engineer the kinetics of SK2, and SK3 channels, we followed the methodology of Mousa ^32^, one S and one F cell models were developed, based on the electrical recordings obtained from S and F MNs, respectively. The anatomy of S and F cell models were developed using reconstructed 3D morphology of spinal mouse MNs ^33,34^. The passive and active properties of the two cell models were optimized to match their respective types from the mouse recordings. Briefly, The S-type and the F-type MNs’ specific membrane resistance (R_m_) for soma, axon hillock, and initial segment were set to 1629 Ω. *cm*^2^ and 738 Ω. *cm*^2^, respectively, while the dendritic membrane resistance was set to a 20 times higher R_m_ following the methodology of Fleshman, et al. ^9^. The specific membrane capacitance (*C*_*m*_) was set to 1μF/cm^2^, and the specific axial resistance (*R*_*a*_) was set to 70 *Ω*. *cm*. fast sodium (Naf), delayed rectifier potassium (Kdr), N-type Ca^2+^ (CaN), and Ca^2+^-activated potassium K(Ca) channels were added to the soma. The initial segment and axon hillock included: Naf, Kdr, and persistent sodium (Nap) channels. The Naf and Kdr channels are responsible for the generation of AP spikes, while CaN channels produce the after-depolarization (ADP) that activates the Ca^2+^-activated potassium K(Ca) channels, which in turn mediate the afterhyperpolarization (AHP). Two types of K(Ca) channels were included in the current computational study: SK2 and SK3, with the goal to reverse engineer their kinetics. A complete list of the passive, and active channel conductances is provided in table S2.

The SK channels were modeled as conductance-based models following Hodgkin and Huxley channel style (equation 1). The channel conductance was modeled as Hill equation, that varies the steady state conductance based on the intracellular Ca concentration (equation 2). The real-time normalized channel conductance (s) was computed using ordinary differential equation of first order with time constant (*τ*_*S*_) (equation 3). The intracellular the intracellular Ca^2+^ dynamics were the same in both the S-type and the F-type MNs following of the methods of Mousa ^32^.

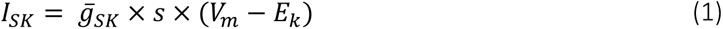

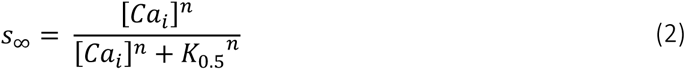

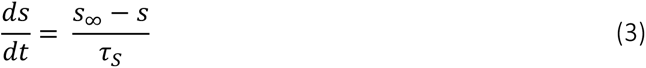

Using the AHP records of the S-type, and F-type MNs, the kinetics of the SK3 and SK2 channels were explored to solve for the 1) the Hill coefficient (n) that represents activation slope of the channel with respect to the Ca^2+^ concentration, 2) Time constant (*τ*_*S*_) that represent activation speed of the channels and 3) the half maximal effective concentration of the intracellular Ca^2+^ *K*_0.5_ that represents the overall channel sensitivity to intracellular Ca^2+^.

#### Statistics

SPSS Software (RRID:SCR_002865) was used to complete all immunohistochemistry and electrophysiology statistical analysis. A two or one way ANOVA was performed on the LCA and density data with Tukey’s post-hoc analysis, A power analysis was completed for all LCA and density measures to determine the appropriate number of cells and animals for analysis. Independent unpaired t-tests were performed for comparisons between SOD and WT groups in density. Alpha was set at a p= 0.05 for significance. All histograms and graphs were created using GraphPad Prism (RRID:SCR_002798). The electrophysiology and computational modeling graphs were all created using Python (RRID:SCR_008394).

## Ethics declaration

Large Language Models (LLMs) were used solely to refine the text of the initial manuscript draft prepared by the authors.

## Data availability statement

All data supporting the results are included in the manuscript figures in addition as accompanied raw data in the supplementary material.

## Acknowledgements

This work was funded by the National Institute of Neurological Disorders and Stroke (award numbers: NS091836 and NS131816) of the National Institutes of Health. The authors would like to thank the following people for their support and contribution to the collection of data for this work: Cierra Ellington, Weston Gelford, Brielle Gatrell, Leila Jaber and Natalie Ehret.

## Disclaimer

The content is solely the responsibility of the authors and does not necessarily represent the official views of the National Institutes of Health.

## References

1 McPhedran, A. M., Wuerker, R. B. & Henneman, E. Properties of Motor Units in a Heterogeneous Pale Muscle (M. Gastrocnemius) of the Cat. J Neurophysiol 28, 85–99 (1965). 10.1152/jn.1965.28.1.85

2 Edström, L. & Kugelberg, E. Histochemical composition, distribution of fibres and fatiguability of single motor units. Anterior tibial muscle of the rat. J Neurol Neurosurg Psychiatry 31, 424–433 (1968).

3 Henneman, E., Somjen, G. & Carpenter, D. O. Functional Significance of Cell Size in Spinal Motoneurons. J Neurophysiol 28, 560–580 (1965).

4 Somjen, G., Carpenter, D. O. & Henneman, E. Responses of motoneurons of different sizes to graded stimulation of supraspinal centers of the brain. J Neurophysiol 28, 958–965 (1965). 10.1152/jn.1965.28.5.958

5 Henneman, E. Relation between size of neurons and their susceptibility to discharge. Science 126, 1345–1347 (1957).

6 Burke, R. E., Levine, D. N., Tsairis, P. & Zajac, F. E., 3rd. Physiological types and histochemical profiles in motor units of the cat gastrocnemius. J Physiol 234, 723–748 (1973).

7 Zengel, J., Reid, S., Sypert, G. & Munson, J. Membrane electrical properties and prediction of motor-unit type of medial gastrocnemius motoneurons in the cat. J Neurophysiol. 53, 1323–1344 (1985).

8 Fleshman, J., Segev, I. & Burke, R. Electrotonic architecture of type-identified alpha-motoneurons in the cat spinal cord. J Neurophysiol 60, 60–85 (1988).

9 Fleshman, J. W., Segev, I. & Burke, R. B. Electrotonic architecture of type-identified alpha-motoneurons in the cat spinal cord. J Neurophysiol 60, 60–85 (1988). 10.1152/jn.1988.60.1.60

10 Deardorff, A. S., et al. Expression of postsynaptic Ca2+-activated K+ (SK) channels at C-bouton synapses in mammalian lumbar -motoneurons. J Physiol 591, 875–897 (2013).

11 Deardorff, A. S., Romer, S. H., Sonner, P. M. & Fyffe, R. E. Swimming against the tide: investigations of the C-bouton synapse. Front Neural Circuits 8, 106 (2014).

12 Gardiner, P. F. Physiological properties of motoneurons innervating different muscle unit types in rat gastrocnemius. J Neurophysiol 69, 1160–1170 (1993). 10.1152/jn.1993.69.4.1160

13 Manuel, M., Meunier, C., Donnet, M. & Zytnicki, D. How much afterhyperpolarization conductance is recruited by an action potential? A dynamic-clamp study in cat lumbar motoneurons. J Neurosci 25, 8917–8923 (2005). 10.1523/JNEUROSCI.2154-05.2005

14 Zengel, J. E., Reid, S. A., Sypert, G. W. & Munson, J. B. Membrane electrical properties and prediction of motor-unit type of medial gastrocnemius motoneurons in the cat. J Neurophysiol 53, 1323–1344 (1985). 10.1152/jn.1985.53.5.1323

15 Dum, R. P. & Kennedy, T. T. Physiological and histochemical characteristics of motor units in cat tibialis anterior and extensor digitorum longus muscles. J Neurophysiol 43, 1615–1630 (1980). 10.1152/jn.1980.43.6.1615

16 Romanul, F. C. DISTRIBUTION OF CAPILLARIES IN RELATION TO OXIDATIVE METABOLISM OF SKELETAL MUSCLE FIBRES. Nature 201, 307–308 (1964). 10.1038/201307a0

17 Alkaslasi, M. R. et al. Single nucleus RNA-sequencing defines unexpected diversity of cholinergic neuron types in the adult mouse spinal cord. Nat Commun 12, 2471 (2021).

18 Blum, J. A. et al. Single-cell transcriptomic analysis of the adult mouse spinal cord reveals molecular diversity of autonomic and skeletal motor neurons. Nat Neurosci 24, 572–583 (2021). 10.1038/s41593-020-00795-0

19 Kaplan, A. et al. Neuronal matrix metalloproteinase-9 is a determinant of selective neurodegeneration. Neuron 81, 333–348 (2014).

20 Misawa, H. et al. Osteopontin is an alpha motor neuron marker in the mouse spinal cord. J Neurosci Res 90, 732–742 (2012). 10.1002/jnr.22813

21 Yamamoto, T. et al. SPP1 expression in spinal motor neurons of the macaque monkey. Neurosci Res 69, 81–86 (2011). 10.1016/j.neures.2010.09.010

22 Chakkalakal, J. V., Nishimune, H., Ruas, J. L., Spiegelman, B. M. & Sanes, J. R. Retrograde influence of muscle fibers on their innervation revealed by a novel marker for slow motoneurons. Development 137, 3489–3499 (2010). 10.1242/dev.053348

23 Friese, A. et al. Gamma and alpha motor neurons distinguished by expression of transcription factor Err3. Proc Natl Acad Sci U S A 106, 13588–13593 (2009). 10.1073/pnas.0906809106

24 Yasvoina, M. V. et al. eGFP expression under UCHL1 promoter genetically labels corticospinal motor neurons and a subpopulation of degeneration-resistant spinal motor neurons in an ALS mouse model. J Neurosci 33, 7890–7904 (2013). 10.1523/JNEUROSCI.2787-12.2013

25 Enjin, A. et al. Identification of novel spinal cholinergic genetic subtypes disclose Chodl and Pitx2 as markers for fast motor neurons and partition cells. J Comp Neurol 518, 2284–2304 (2010). 10.1002/cne.22332

26 Ringer, C., Weihe, E. & Schutz, B. Calcitonin gene-related peptide expression levels predict motor neuron vulnerability in the superoxide dismutase 1-G93A mouse model of amyotrophic lateral sclerosis. Neurobiol Dis 45, 547–554 (2012). 10.1016/j.nbd.2011.09.011

27 Muller, D. et al. Dlk1 promotes a fast motor neuron biophysical signature required for peak force execution. Science 343, 1264–1266 (2014). 10.1126/science.1246448

28 McCombe, P. A. & Henderson, R. D. Effects of gender in amyotrophic lateral sclerosis. Gend Med 7, 557–570 (2010). 10.1016/j.genm.2010.11.010

29 Ishihara, A., Nagatomo, F., Fujino, H., Kondo, H. & Ohira, Y. Decreased succinate dehydrogenase activity of gamma and alpha motoneurons in mouse spinal cords following 13 weeks of exposure to microgravity. Neurochem Res 38, 2160–2167 (2013). 10.1007/s11064-013-1124-y

30 Mahrous, A. A., Mousa, M. H. & Elbasiouny, S. M. The Mechanistic Basis for Successful Spinal Cord Stimulation to Generate Steady Motor Outputs. Front Cell Neurosci 13, 359 (2019).

31 Deutsch, A. J. & Elbasiouny, S. M. Dysregulation of persistent inward and outward currents in spinal motoneurons of symptomatic SOD1-G93A mice. J Physiol 602, 3715–3736 (2024). 10.1113/JP286032

32 Mousa, M. H. Reverse Engineering Spinal Motoneuron Properties In Slice And Whole-Cord Preparations Using Computational Modeling, Wright State University, (2022).

33 Li, Y., Brewer, D., Burke, R. E. & Ascoli, G. A. Developmental changes in spinal motoneuron dendrites in neonatal mice. The Journal of Comparative Neurology 483, 304–317 (2005).

34 Ascoli, G. A., Donohue, D. E. & Halavi, M. NeuroMorpho.Org: A Central Resource for Neuronal Morphologies. Journal of Neuroscience 27, 9247–9251 (2007). 10.1523/JNEUROSCI.2055-07.2007

