## Supplemental figures for "Circular continuum of alpha motoneuron types"

**Supporting Information for**
### Circular continuum of alpha motoneuron types

Teresa L. Garrett[1], Lee Wintermute[2], Maggie Armitage[1], Shelby Ward[2], Ibrahim Abdul Halim[2], Mohamed H. Mousa[1], Kalin Gerber[2], and Sherif M. Elbasiouny[1,2]

Sherif M. Elbasiouny, PhD


**This PDF file includes:**

Figures S1 to S2
Tables S1 to S2

**Data sets included as supporting information:**

Protocol 1 and 2 data located: OPN_MMP9 data.xlsx

Protocol 3 data located: SK2_SK3labeling.xlsx

Protocol 3 data located: opn_sk2_3 data.xlsx

Electrophysiology data located: electrophysiology data.xlsx

Supplemental data figure 1 located: C57 M_F data.xlsx

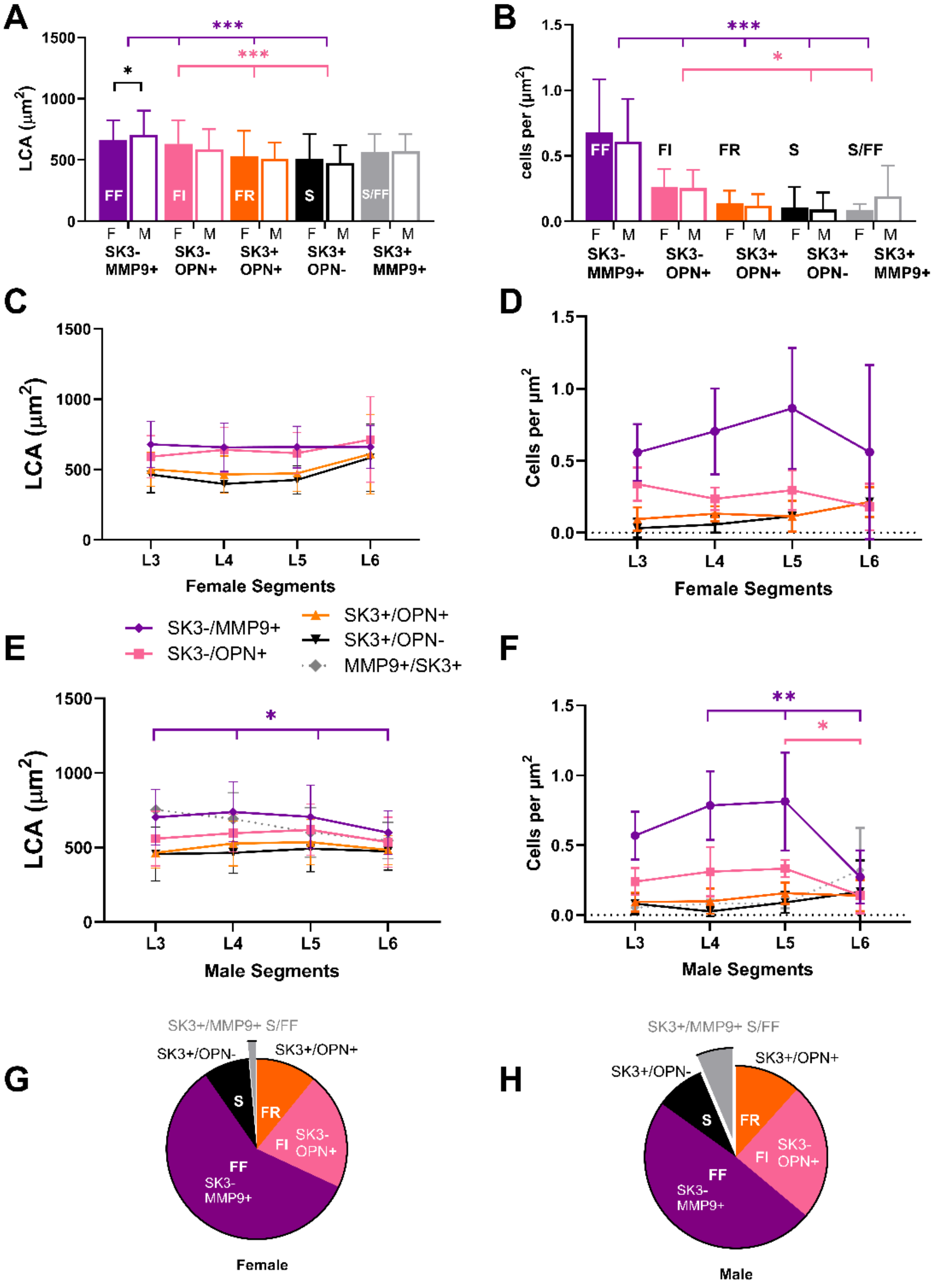


**Figure S1.** Data of α-MN types in the lumbar spinal cord of adult female and male non-transgenic C57BL/6J mice. Largest cross-sectional area (LCA, A) (N of each type female: FF=396, FI=151, FR=82=, S=60, and S/FF=10, male: FF=390, FI=174, FR=86=, S=60, and S/FF=51)) and density (B) of α-MN types in female (marked as F) and male (marked as M) mice. Distribution of α-MN types' LCA across lumbar segments in female (C) and male (E) mice. Density of α-MN types across lumbar segments in female (D) and male (F) mice. Percentage composition of α-MN types in the adult lumbar spinal cord of female (G) and male (H) mice.

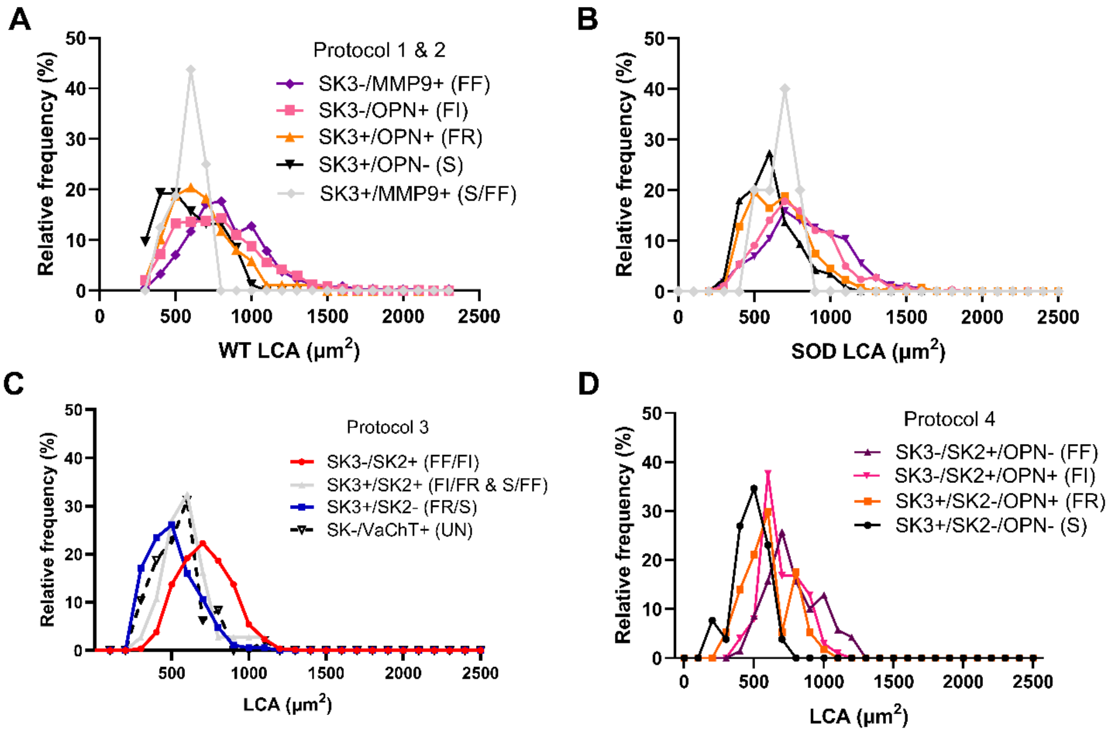


**Figure S2.** LCA frequency distributions in SOD1-G93A WT mice for each α-MN type for each labeling protocol showing significant overlap of size and cell type in all labeling protocols.

| Section | Parameter | Value | |
|---|---|---|---|
| | | S-type MN (SK3) | F-type MN (SK2) |
| **Soma** | $R_m$, $\Omega.cm^2$ | 1629 | 738 |
| | $\bar{g}_{Naf}$, $S\,cm^{-2}$ | 0.06 | 0.075 |
| | $\bar{g}_{Kdr}$, $S\,cm^{-2}$ | 0.34 | 0.26 |
| | $\bar{g}_{CaN}$, $S\,cm^{-2}$ | 0.2 | 0.04 |
| | $\bar{g}_{SK2}$, $S\,cm^{-2}$ | 0 | 0.011 |
| | $\bar{g}_{SK3}$, $S\,cm^{-2}$ | 0.0054 | 0 |
| **Axon Hillock and initial segment** | $R_m$, $\Omega.cm^2$ | 1629 | 738 |
| | $\bar{g}_{Naf}$, $S\,cm^{-2}$ | 0.45 | 0.45 |
| | $\bar{g}_{Kdr}$, $S\,cm^{-2}$ | 0.2 | 0.14722 |
| | $\bar{g}_{NaP}$, $S\,cm^{-2}$ | 0.0004 | 0.0004 |
| **Dendrites** | $R_m$, $\Omega.cm^2$ | 32,580 | 14,760 |

**Table S1.** Ion Channel Conductance in Various Structures of F and S MN Computer Models.

| Segment | | FF | FI | FR | S | S/FF |
|---|---|---|---|---|---|---|
| L3 | WT<br>SOD | 688.62±219.73 (N=184)<br>724.36±279.92 (N=173) | 630.29±209.32 (N=173)<br>688.34 317.11 (N=92) | 561.48±116.42 (N=28)<br>526.67±109.43 (N=21) | 573.64±173.17 (N=35)<br>549.0±155.57 (N=38) | 375.94 19.85 (N=2)<br>618.61 335.13 (N=5) |
| L4 | WT<br>SOD | 805.43±248.48 (N=277)<br>856.84±214.51 (N=188) | 775.24±263.94 (N=258)<br>763.53 208.13 (N=167) | 649.61±209.4 (N=47)<br>671.9±200.85 (N=30) | 469.64±148.63 (N=16)<br>583.72±146.38 (N=31) | N=0<br>N=0 |
| L5 | WT<br>SOD | 941.66±297.03 (N=363)<br>878.85±264.5 (N=162) | 802.68±261.38 (N=319)<br>834.07 263.83 (N=154) | 706.15±239.11 (N=59)<br>714.37±272.25 (N=44) | 625.9±197.78 (N=15)<br>671.73±198.7 (N=13) | N=0<br>645.59 188.57 (N=3) |
| L6 | WT<br>SOD | 863.13±248.7 (N=291)<br>890.87±259.05 (N=154) | 854.5±285.73 (N=216)<br>864.55 278.05 (N=126) | 715.21±202.18 (N=52)<br>691.06±158.79 (N=38) | 647.71±188.67 (N=17)<br>641.14±186.19 (N=35) | 609.21 80.53 (N=14)<br>637.7 220.3 (N=11) |

**Table S2.** Mean LCA of both SOD1-G93A WT and SOD mice across each segment for each α-MN type using protocols 1 and 2.
